# Four decades of plant community change along a continental gradient of warming

**DOI:** 10.1101/313379

**Authors:** Antoine Becker-Scarpitta, Steve Vissault, Mark Vellend

## Abstract

Many studies of individual sites have revealed biotic changes consistent with climate warming (e.g., upward elevational distribution shifts), but our understanding of the tremendous variation among studies in the magnitude of such biotic changes is minimal. In this study we re-surveyed forest vegetation plots 40 years after the initial surveys in three protected areas along a west-to-east gradient of increasingly steep recent warming trends in eastern Canada (Québec). Consistent with the hypothesis that climate warming has been an important driver of vegetation change, we found an increasing magnitude of changes in species richness and composition from west to east among the three parks. For the two mountainous parks, we found no changes in elevational species’ distributions in the eastern most park where warming has been minimal (Forillon Park), and significant upward distribution shifts in the centrally located park where the recent warming trend has been marked (Mont-Mégantic). Community temperature indices (CTI), reflecting the average affinities of locally co-occurring to temperature conditions across their geographic ranges (“species temperature indices”), did not change over time as predicted. However, close examination of the underpinnings of CTI values suggested a high sensitivity to uncertainty in individual species’ temperature indices, and so a potentially limited responsiveness to warming. Overall, by testing *a priori* predictions concerning variation among parks in the direction and magnitude of vegetation changes, we have provided stronger evidence for a link between climate warming and biotic responses than otherwise possible, and provided a potential explanation for large variation among studies in warming-related biotic changes.

## Introduction

Climate is a dominant driver of large-scale plant distributions (Pearson & Dawson, 2003). On smaller spatial and temporal scales, changes in local climatic conditions can lead to modifications of species’ abundances (Vellend *et al.*, 2017), risks of extinction (Parmesan & Yohe, 2003; Rooney *et al.*, 2004; Urban, 2015), phenology (Menzel *et al.*, 2006; Cleland *et al.*, 2007), distributions (Kelly & Goulden, 2008; Lenoir *et al.*, 2008; Bertrand *et al.*, 2011) and local adaptation (Aitken *et al.*, 2008). Although many such changes have been observed in previous studies, the magnitude of response varies tremendously from study to study, and we have only a limited understanding of the processes underlying this variation.

Most of the world’s natural vegetation is dominated by long-lived perennials plants (Grime, 1977), and so we expect vegetation responses to environmental change to occur slowly relative to the time span of a few years (or less) typical of ecological studies (Tilman, 1989). A key strategy used to assess longer-term temporal changes in plant communities is the resurvey of plots initially surveyed decades ago, often referred to as “legacy” studies (Vellend *et al.*, 2013a; Chytrý *et al.*, 2014; Hédl *et al.*, 2017; Perring *et al.*, 2017). An important limitation of such studies is their constrained ability to test the ecological mechanisms underlying temporal community change. Indeed, most legacy studies pertain to a single site, meaning a set of plots within an area sharing a similar climate and history, in which case community change might be caused by many local changes, such as ongoing land use (Hermy & Verheyen, 2007; Kampichler *et al.*, 2012; Newbold *et al.*, 2015), historical management legacies (Vanhellemont *et al.*, 2014; Becker *et al.*, 2016; Perring *et al.*, 2017), nitrogen deposition (Becker-Scarpitta *et al.*, 2017) or grazing (Frerker *et al.*, 2014; Vild *et al.*, 2016).

Causes of community change at a single site are often assessed by comparing observed changes in community composition across space or time with predictions based on drivers of interest, such as the climate warming. For instance, as predicted by the climate warming hypotheses, many species have experienced a shift in distribution towards higher elevations (Gottfried *et al.*, 2012; Pauli *et al.*, 2012; Stockli *et al.*, 2012; Sproull *et al.*, 2015) or latitudes (Parmesan *et al.*, 1999; Hickling *et al.*, 2006; Boisvert-Marsh *et al.*, 2014; but see VanDerWal *et al.*, 2012). Given that plant species richness tends to be greater in warmer areas, a local-scale increase in richness is also predicted due to warming, at least in the absence of severe moisture stress (Vellend *et al.*, 2017). Finally, if each species is first characterized by its geographic affinity with different temperature conditions (using a “Species Temperature Index”), then the average affinity across species in a local community (the “Community Temperature Index”) is predicted to increase in response to warming (Devictor et al. 2008, 2012). Although there have been considerable advances in testing these predictions in single-site studies (local scale), explicit tests of predictions comparing multiple sites (regional scale) are needed to improve our knowledge and ability to predict biodiversity responses to climate changes (Verheyen *et al.*, 2017).

Here we report analyses of changes in forest plant communities over four decades at three sites strategically chosen to be in areas covering a range of recent climate warming trends in eastern North-America (Québec, Canada). To assess temporal changes, we have revisited sites where botanical legacy data were collected in the 1970s, during the time that many provincial parks were being planned and established in Québec. Plots were widely distributed throughout each park and were typically placed in mature forest stands. Since the time of the original surveys, these forests have not experienced any major anthropogenic disturbances, thus minimizing possible confounding causes of vegetation change.

The province of Québec (Canada) spans >1000 km east-west, over which there is a marked gradient of warming over the past ~60 years (see Appendix S1 and Yagouti *et al.*, 2008). At the tip of the Gaspé Peninsula, the location of our most easterly site, Forillon National Park (Fig. 1), warming has been least pronounced, likely due to the climatic buffering effect of the Atlantic Ocean (see Appendix S1). In contrast, Gatineau Park in continental western Québec has experienced marked warming, with Mont-Mégantic Provincial Park in between both geographically and in terms of the magnitude of warming (Fig. 1 and S1,Yagouti *et al.*, 2008). To the best of our knowledge, no study has used legacy data to specifically test for contrasting vegetation responses in sites with variable warming trends (but see Menzel *et al.*, 2006 for phenological responses to different warming trends).

**Figure 1:**
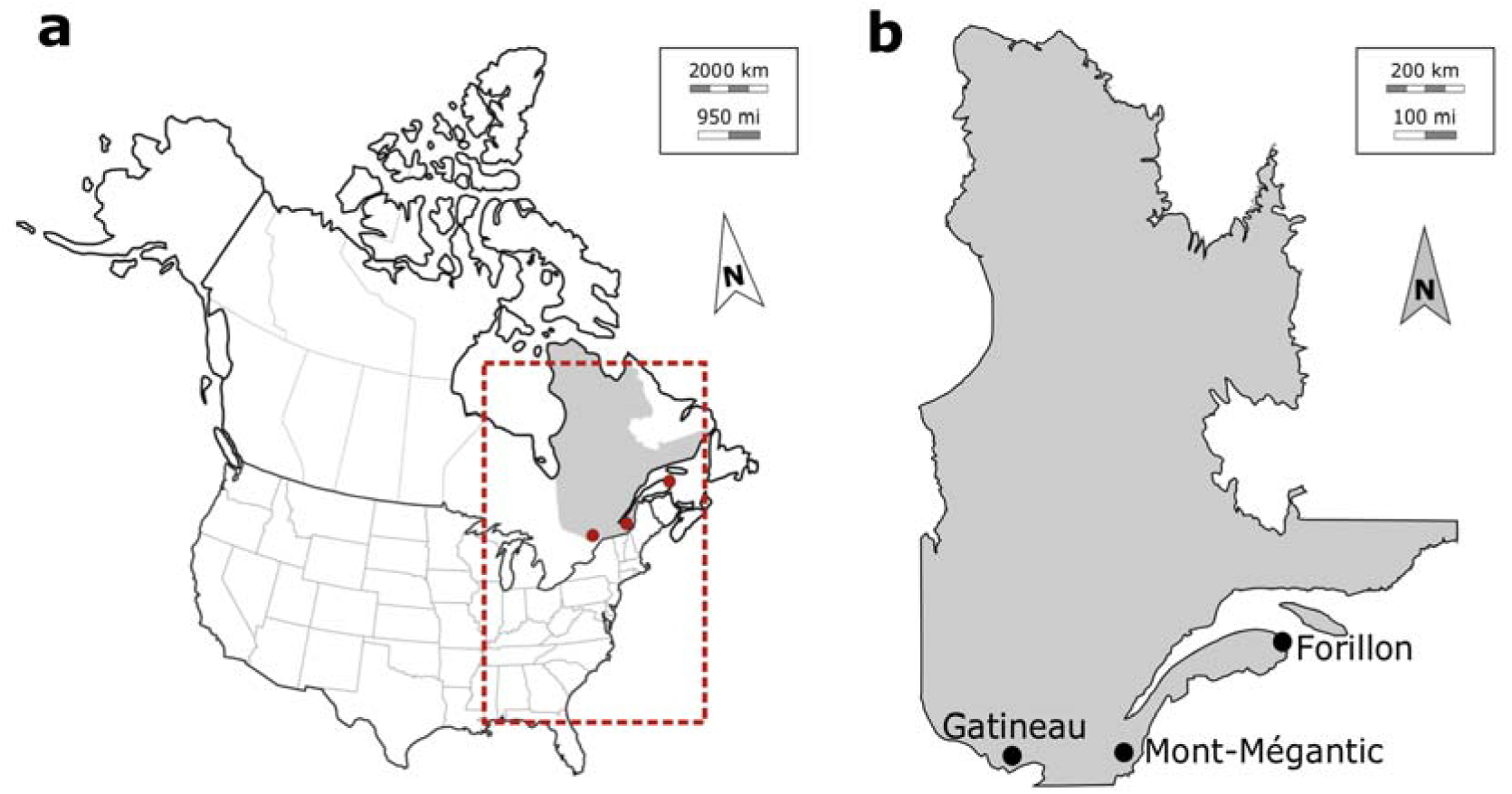
Location of study sites in (a) Canada and (b) the Province of Québec. The red box in (a) shows the area used for extraction of species occurrences in the calculation of Species Temperature Indices (STI): 60°-90°W; 30°- 60°N

Our core hypothesis is that areas with greater warming will have experienced stronger vegetation changes than areas with less warming (Chen *et al.*, 2011; Wang *et al.*, 2017). We take advantage of this unique combination of original studies along a warming gradient to perform a regional-scale analysis of temporal change of forest plant communities. Results for Mont-Mégantic, including significant upward elevational distribution shifts and increased local species richness, were reported in a previous paper (Savage & Vellend, 2015), to which we here add data for Gatineau Park (stronger warming trend) and Forillon Park (weaker warming trend). We tested the following specific predictions: (1) Significant upward elevational distribution shifts have occurred at Mont-Mégantic (already observed) but not at Forillon Park (tested in this paper). (Elevational variation in Gatineau Park is minimal – insufficient to test for temporal shifts in species distributions.) The magnitude of (2) the temporal change in species richness, (3) the temporal change in community composition (R^2^ from the “time” effect in a multivariate analysis), and (4) the temporal change in Community Temperature Index (CTI) vary in magnitude among parks as follows: Forillon < Mont-Mégantic < Gatineau.

## Materials and methods

### Study region and sites

We studied vegetation change in three north-temperate forest sites in eastern Canada (Québec), spanning ~1000 km from Forillon National Park in eastern Québec, to Mont-Mégantic Provincial Park in central Québec and Gatineau National park in the western part of the province (Fig. 1). For all three parks, there has been no logging or forest management during the period of study.

Forillon National Park, located at the eastern extremity of the Gaspé peninsula (48°54 ′N, 64°21 ′W), was created in 1970 and covers 245 km^2^, with our study plots ranging in elevation from ~50 to 500 m a.s.l. The vegetation at Forillon is characterized in large part by boreal species, such as *Abies balsamea* (L.) Mill., *Picea glauca* (Moench) Voss and *Betula papyrifera* Marshall. At low elevation, temperate deciduous or mixed forests are dominated by *Acer saccharum* Marsh. and *Betula alleghaniensis* Britt. (Majcen, 1981).

Mont-Mégantic Provincial Park is located in the Eastern Townships region of Québec (45°27 ′N, 71°9 ′W), about 650 southwest of Forillon Park and 15 km north of the U.S. borders with New Hampshire and Maine. The park was created in 1994 (logging ceased in the 1960s prior to park planning) and covers ~55 km^2^. Our study plots range in elevation between ~460 and 1100 m a.s.l. Vegetation patterns are very similar to Forillon, with a somewhat more visually evident elevational gradient: at low elevations, temperate deciduous forests are dominated by *Acer saccharum* Marsh., *Fagus grandifolia* Ehrh. and *Betula alleghaniensis* Britt., while at high elevation boreal forests are composed largely of *Abies balsamea* (L.) Mill. and *Picea rubens* Sar. (Marcotte & Grandtner, 1974).

Gatineau Park is located in southwestern Québec (45°35 ′N 76°00 ′W), in the Outaouais region, 360 km west of Mont-Mégantic. The park was established in 1938, covers 361 km^2^, with relatively little elevational variation compared to the other parks (250 m elevational range). Contrary to Forillon and Mont-Mégantic, our vegetation sampling was not spread throughout the entire park (access to certain sectors of the part is restricted). Our study area (~30 km^2^) is largely dominated by *Acer saccharum* Marsh and *Fagus grandifolia* Ehrh., with a few more southerly tree species such as *Tilia americana* L.*, Quercus rubra* L.*, Quercus alba* L. *or Fraxinus americana* L. as well.

### Data set

All original vegetation surveys were conducted using phytosociological methods (Marcotte & Grandtner, 1974; Chartrand, 1976; Majcen, 1981). In fixed-area plots (see below), authors made a full list of vascular plant species in different strata (i.e. canopy trees, shrubs, herbs) with abundance coefficients per species assigned following the scale of Braun-Blanquet *et al.* (1952). In our analyses, we pooled shrubs and herbs into a single “understorey” stratum and given the limited representation of the tree community in smaller plots (90m^2^, see below), we focused all analyses on the understorey data. For analyses, Braun-Blanquet classes were converted to a percentage value representing the mid-point of a given abundance class.

None of the original survey plots were permanently marked, but for all three parks plot coordinates were reported in maps and/or tables. As such, plots are considered “semi-permanent”, which introduces the possibility of pseudo-turnover due to relocation uncertainty (Stockli *et al.*, 2012; Vellend *et al.*, 2013a; Hédl *et al.*, 2017; Kapfer *et al.*, 2017). However, previous studies have shown that conclusions are robust to uncertainty in plot relocation, which adds statistical noise but not systematic bias (Kopecký & Macek, 2015). In our study, original surveyors tended to sample mature forest stands where spatial heterogeneity was relatively low, thus reducing any effects of plot relocation uncertainty. We used original plot maps and environmental descriptions (elevation, slope, aspect) to select potential locations for resurvey plots in a GIS (QGIS Development Team 2016, Open Source Geospatial Foundation Project). Potential locations were visited in the field, with the final location of a given plot determined by the best match to the original location and description. Logistical limitations prevented us from resurveying all original plots in Forillon and Gatineau. At Mont-Mégantic, all plots within the current park boundary were surveyed in 2012 (see Savage & Vellend, 2015). Plot selection for our recent surveys followed several criteria: (i) plots occurred in forest, excluding swamps or bogs; (ii) plots were accessible via <3-4 hours hiking off of trails (abandonment of old forest roads and trails since the 1970s has reduced accessibility); (iii) plots had not obviously experienced recent major natural disturbances (e.g., storms, fire, or insect outbreaks); (iv) in the original survey the plots were sampled in mature stands that have since maintained forest cover (i.e., no early successional dynamics in the intervening period).

At Forillon, the original survey was conducted in June-September 1972 in 256 vegetation plots of 500 m^2^ distributed throughout the park (Majcen, 1981). We resurveyed 49 plots during July and August of 2015. At Mont-Mégantic, the vegetation was originally surveyed in 1970 in 94 plots, almost half of which were outside of the current park boundaries. The plot size was 400 m^2^ in coniferous forest and 800 m^2^ in broadleaved forests (Marcotte & Grandtner, 1974). Among the 94 original plots, 48 were revisited within the current park limits at Mont-Mégantic in 2012, with results reported in Savage & Vellend (2015). In Gatineau Park, surveys were conducted in 1973 in 33 plots of 90 m^2^ during the summer in 1973 (Chartrand, 1976) and 28 plots were resurveyed in summer 2016. We harmonized taxonomy across all three parks and two time periods (see below), so the Mont-Mégantic data are not precisely the same as reported in Savage & Vellend (2015). The study design was perfectly balanced within parks for statistical analysis (i.e., the same number of plots in the original and recent surveys).

### Taxonomy

Our taxonomical reference for vascular plants was the Taxonomic Name Resolution Service v4.0 (assessed in Feb 2017: http://tnrs.iplantcollaborative.org).

Our data set was collected by five different survey teams, one for each of the three original surveys: Forillon: Majcen (1981); Mont-Mégantic: Marcotte & Grandtner (1974); Gatineau:Chartrand (1976), one for the recent Mont-Mégantic survey: Savage & Vellend (2015), and one for the recent Forillon and Gatineau surveys (A. Becker-Scarpitta and assistants). Most plants were identified to the species level in the same way across surveys, such that the only harmonization step for these taxa was to standardize names, which may have changed over time. In many cases, however, coarser levels of taxonomic resolution (e.g., a pair of similar species not identified to the species level) were used in some but not all surveys, or the timing of different surveys created doubt about the likelihood of comparable detection abilities (e.g., for spring ephemeral plants) (see Appendix S2 for details on taxonomic standardization). In these cases, the coarser level of resolution was applied to all data sets, or species were removed to maximize comparability. We deposited all specimens identified at the species level to the Marie-Victorin herbarium (Institut de Recherche en Biologie Végétale, Université de Montréal, Canada) and all locations were entered into the GBIF database (GBIF - https://www.gbif.org).

### Community Temperature Index (CTI)

A predicted response of communities to warming is a temporal increase in the Community Temperature Index (CTI), which we calculated for all plots in each survey. CTI was calculated as the abundance-weighted average of the Species Temperature Index (STI) across all species in a given plot. The STI for a given species is the median of the long-term (1960-2010) mean annual temperatures calculated across all known occurrences of the species (Devictor *et al.*, 2008). To calculate STIs, we compiled an independent dataset by extracting all recorded occurrences for each species in the Botanical Information and Ecology Network (BIEN - http://bien.nceas.ucsb.edu/bien/; Enquist *et al.*, 2016) in eastern North America: 60° to 90°W; 30° to 60°N (red box of Figure 1). We excluded occurrences further west, in order to control the range of variation in precipitation (precipitation decreases markedly to the west of the deciduous forest biome). Our STIs thus reflect temperature affinities under precipitation conditions most comparable to those found in our study region. For each occurrence point, we extracted the annual mean temperature from ANUSPLIN, a model developed by Natural Resources Canada (http://cfs.nrcan.gc.ca/projects/3; McKenney et al. 2006). The abundance-weighted version of CTIw was calculated for each plot *j* as:

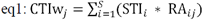

The STI of species *i* is weighted by the relative abundance (RA) of species *i* in plot *j* (RA = the species local abundance divided by the sum of all S species’ abundances in that plot). Given some surprising results concerning CTIw, we also explored analyses of the unweighted version, CTIuw (median STI across species with no weighting for abundance), thus focusing on which species were present in a given plot rather than their relative abundances.

STI values were calculated only for species identified at the species level and with more than 50 occurrences in the BIEN database (see Appendix S3 – Species Temperature Index database). Note that compared to Savage & Vellend (2015) we used improved climate data (ANUSPLIN instead of WORLDCLIM) and updated distribution data (BIEN instead of GBIF), thus leading to the potential for different results.

### Statistical analysis

All statistical analyses were performed in R v.3.4.2 (R Foundation for Statistical Computing 2017). To test for upward elevational shifts in species distributions at Forillon and Mont-Mégantic, we selected species occurring in at least four plots per survey in a given park. For each species in each park we calculated the average abundance-weighted elevation across occurrences. We then conducted linear mixed effect models (LMM, function *lmer*, package ‘lme4’ v.1.1-14,Bates *et al.*, 2015) testing for a fixed effect of time period on abundance-weighted mean elevation, with species as a random effect to account for the paired sampling structure of the data (each species observed in each time period).

We first studied the relationship between α-diversity (species richness) and time using LMMs including time, elevation and the time*elevation interaction (if significant) as fixed effects, and plot ID as a random effect. Because Gatineau has a negligible elevation gradient, we used a model for this park with only time and plot ID as a random effect. Coefficients of determination were expressed as marginal R^2^ (R^2^_m_) and conditional R^2^ (R^2^_c_) using the function *r.squaredGLMM*, package ‘MuMIn’ v.1.40.0 (Nakagawa & Schielzeth, 2013).

We then explored temporal change in β-diversity (i.e. the variability in species composition among communities) using permutational analysis of multivariate dispersion (PERMDISP). This analysis assessed the multivariate homogeneity of group dispersions based on Bray-Curtis distances (also called percentage-difference distance), with significance testing via permutation (function *betadisper*, package ‘vegan’ v.2.4-4,Anderson *et al.*, 2006). A decrease in the multivariate distance between plots and the time-specific centroid is interpreted as biotic homogenization, while an increase indicates biotic differentiation.

To examine changes in community composition over time, we used permutational analysis of variance (PERMANOVA, with Bray-Curtis distances) using 999 permutations (function *adonis*, package ‘vegan’) (Anderson, 2001). We used the R^2^ values from the PERMANOVA models as quantification of the magnitude of temporal change in order to compare among parks. We used non-metric multidimensional scaling (NMDS) with Bray-Curtis distances for visualization (function *metaMDS*, package ‘vegan’).

Temporal changes in the Community Temperature Index (CTI) were tested using LMMs for both weighted and unweighted versions of CTI (CTIw and CTIuw, respectively). Model structure was identical to the model for species richness. We included the interaction between time and elevation only if significant.

## Results

### Species elevational distributions

In Forillon, where there has been the least warming in recent decades, there was no significant temporal change, on average, in understorey species’ elevational distributions (original survey mean = 195.4 ± 12.3 (SE) m; recent = 206.8 ± 12.3 m, t = 0.85, p = 0.41, Fig. 2a, see Appendix S4 for species-by-species data). In contrast, a significant upward elevational shift was observed at Mont-Mégantic, which has experience marked warming (original mean = 622.1 ± 10 m, recent mean = 660.94 ± 10 m, t = 4.67, p < 0.001, Fig. 2b). At Mont-Mégantic, on average species’ distributions have shifted 39 m towards higher elevations (~10 m.decade^−1^), and this was consistent along the spatial gradient (Fig. 2b; see also Savage & Vellend 2015).

**Figure 2.**
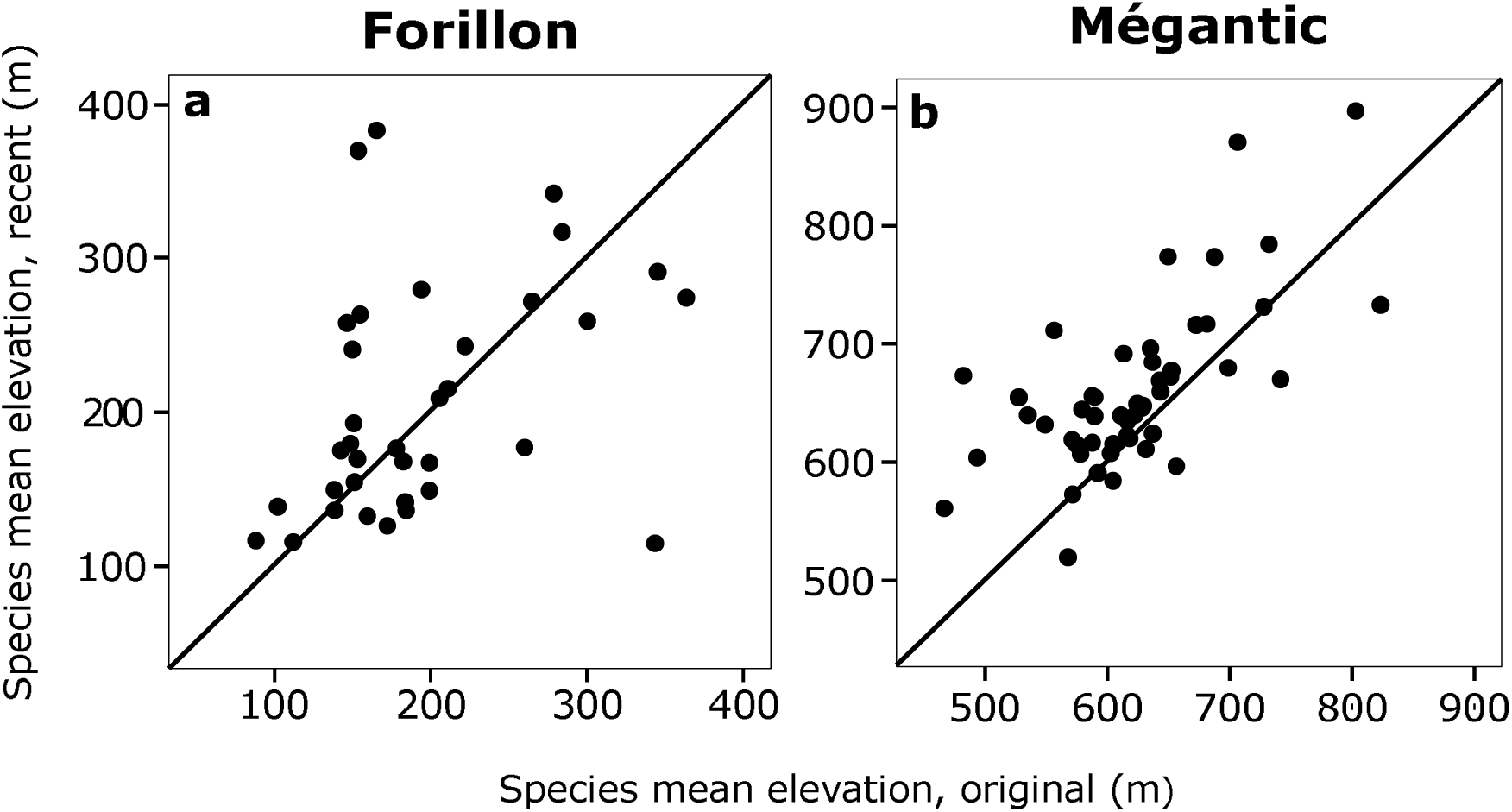
Changes over time in species’ elevational distributions at (a) Forillon, n=35 species, F=0.70, p=0.41 – no significant shift in elevation, and (b) Mont Mégantic, n=50 species, F=22.72, p<0.001 – significant upward shift in elevation. The diagonal line (1:1) represents no elevational change over time. Each point represents one species (occurring in minimum four plots per survey); see Appendix S4 for data.

### Species richness

At Forillon, for plot-level species richness (α-diversity) we found no significant temporal change (Table 1 and 2), and the weak negative trend of richness with elevation was not significant (Fig 3d, Table 1). Across all plots we observed 18 fewer understorey species in the recent survey (65 species) than in the original survey (83 species); 27 species present in original survey were not found in the recent one, while we found 9 new species (Table 2). It is important to note that these are not likely to be gains and losses to and from the entire park, but only to and from this set of semi-permanent plots.

**Figure 0.**
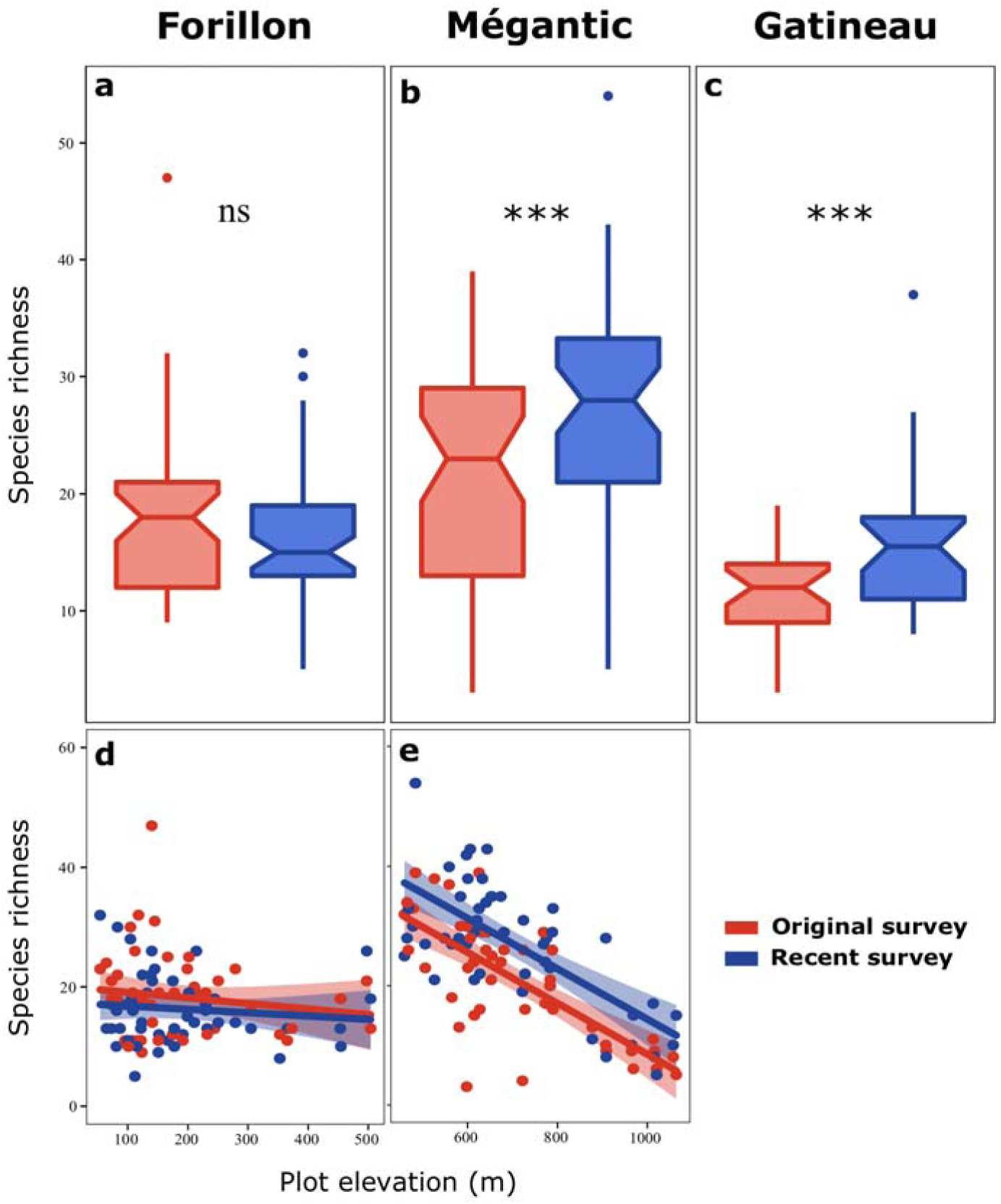
Temporal changes in understorey species richness. (a-c) Box plots of original and recent species richness per plot in the three parks. (d-e) Linear relationships between species richness and elevation in the original and recent surveys at Forillon (n=49*2 plots, no significant relationship for either original or recent surveys, see Table 1), and Mont-Mégantic (n=48*2 plots, significant relationship for both original and recent surveys, see Table 1). The colored polygons around each regression line represent 95% confidence intervals. *** p<0.001

**Table 1.**
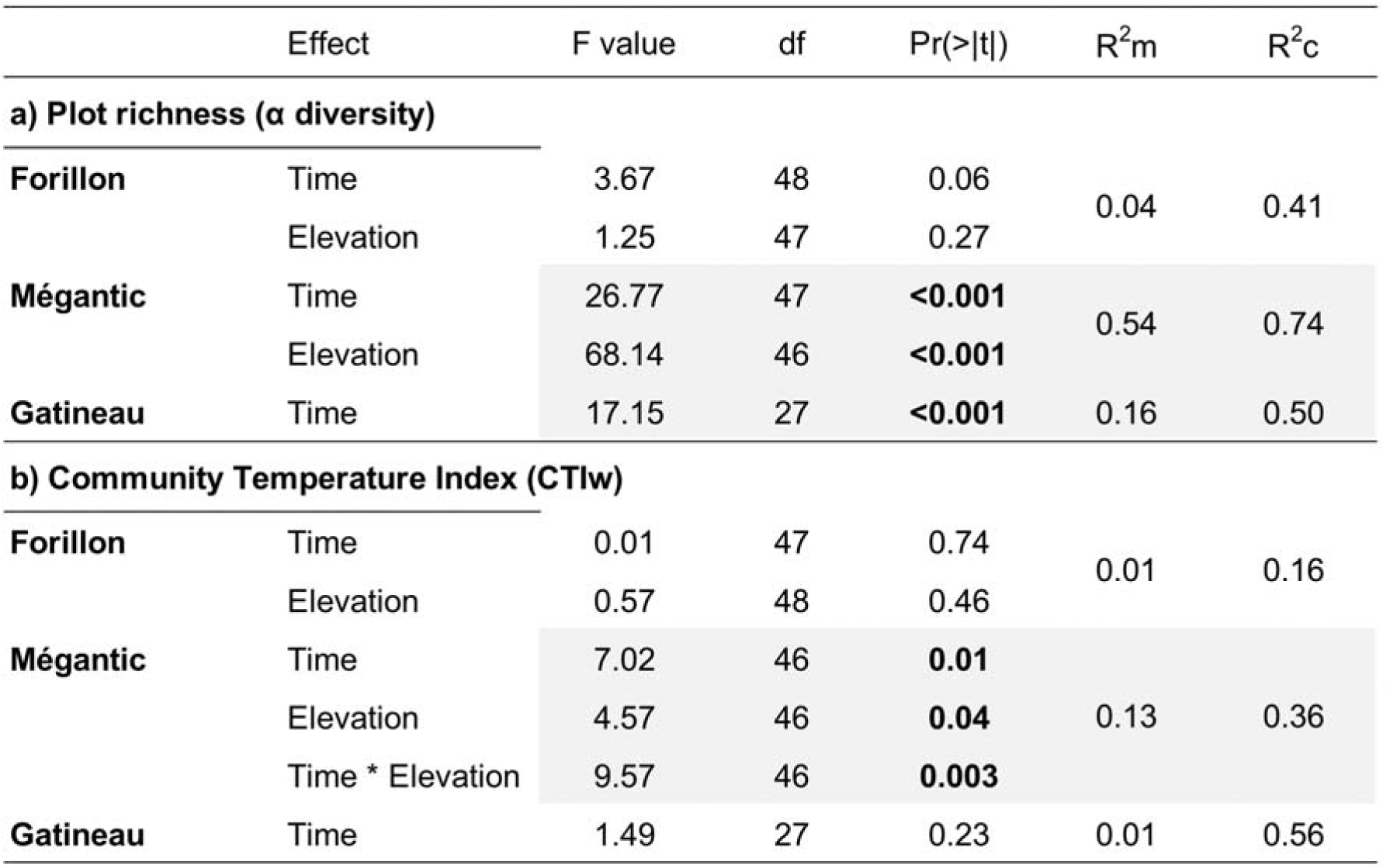
Results of linear mixed models (LMMs) predicting species richness and community temperature indices (CTIw). R^2^m is the marginal R^2^, measuring the proportion of variance explained by fixed effects; R^2^c is the conditional R^2^, giving the proportion of variance explained by both fixed and random effects.

**Table 2.** Temporal changes in total species numbers and plot-level species richness (α- diversity). The total number of species observed across all plots is broken down into those shared, lost, or gained between the original and recent surveys. For plot-level richness, means ± SE are reported. Shading indicates significant statistical differences (p < 0.05, see Table 1 for statistical tests)

| | Total species number | | | | | α-diversity | |
| --- | --- | --- | --- | --- | --- | --- | --- |
|  | Original | Recent | Shared | Losted | Gained | Original | Recent |
| **Forillon** | 83 | 65 | 56 | 27 | 9 | 18.2 ± 1 | 16.4 ± 0.8 |
| **Mégantic** | 87 | 92 | 79 | 8 | 13 | 21.2 ± 1.5 | 27.0 ± 1.5 |
| **Gatineau** | 70 | 90 | 58 | 12 | 32 | 11.6 ± 0.8 | 15.9 ± 0.8 |

At Mont-Mégantic richness declined significantly with elevation in both time periods (original: t = −6.97, p < 0.001; recent: t = −6.91, p < 0.001, Fig. 3e and Table 1). Similar numbers of understorey plant species overall were found in the recent and original surveys (92 and 87 species, respectively); 8 species from the original survey were not found, while we recorded 13 new species in recent survey (Table 2). Mont-Mégantic showed a significant increase over time in the plot-level richness of understorey species (27% increase on average, see Fig. 3b and Tables 1 and 2), and this increase was consistent across the elevational gradient (Fig. 3e).

Finally, in Gatineau Park, plot-level species richness increased significantly by an average of 38% (t = 4.14, p < 0.001, Fig. 3c, Table 1 and 2). Overall, we found 20 more species in the recent survey than in the original survey. Gatineau showed the largest study-wide gain in species, with 32 new species observed in the recent survey and 12 species from the original survey not observed in recent one (Table 2).

### Community composition and heterogeneity

At none of the three sites was there significant temporal change in β-diversity (Table 3). However, we observed highly significant shifts in understorey community composition for all study sites (Table 3), although in the two-dimensional NMDS ordinations the shifts appear fairly subtle (Fig. 4). The magnitude of the understorey compositional shifts (R^2^) increased from Forillon (5%) to Mont-Mégantic (8%) to Gatineau (10%) (Table 3). Appendix S5 reports the list of species frequencies and Appendix S6 shows an NMDS ordination of all plots from all parks together.

**Figure 4.**
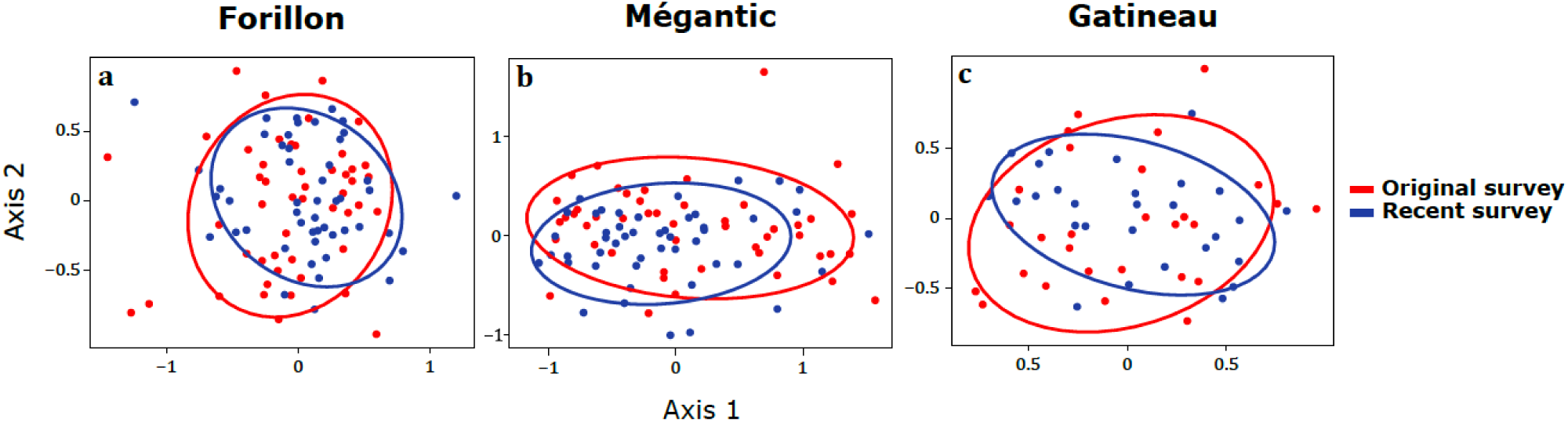
Non-metric multidimensional scaling (NMDS) ordinations of understorey communities across time for (a) Forillon, stress = 0.94; (b) Mont-Mégantic, stress=0.97 and (c) Gatineau, stress=0.97. Each point represents a survey plot, and colors refer to the time-period of surveys (red: original survey; blue: recent survey). Ellipses show 75% confidence limits for each time-period. We used two dimensions and Bray-Curtis distances. For a single ordination with species names see Appendix S6.

**Table 3.**
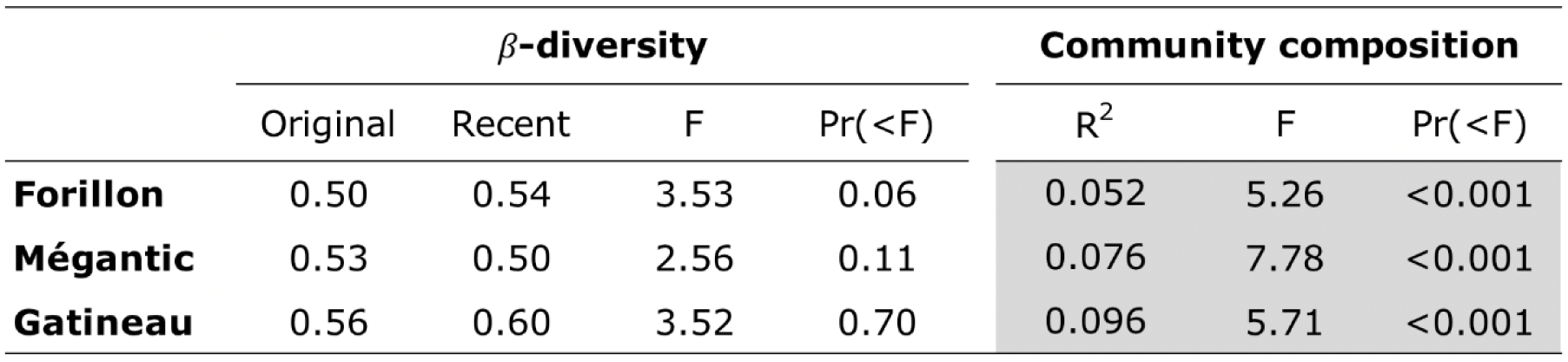
Tests for temporal shifts in β-diversity (PERMDISP) and community composition (PERMANOVA) of understorey communities between original and recent surveys. β-diversity is the mean distance between each plot and the time-specific centroid in multivariate space (Bray-Curtis distances). R^2^ is the proportion of variation in community composition explained by time. Statistical significance levels were calculated with 999 permutations.

### Community temperature indices (CTI)

The only significant temporal change in Community Temperature Indices (CTIw) was found at Mont-Mégantic, and the change was negative, the opposite of the predicted direction. We detected no significant changes in CTI in Forillon or Gatineau (Fig. 5 and Table 1). At Forillon, there was no significant relationship between CTI and elevation for either the original or recent survey (Fig. 5d and Table 1), nor was there any relationship for the original survey at Mont-Mégantic (Fig. 5e and Table 1). For the recent survey at Mont-Mégantic, there was a significant negative relationship between CTIw and elevation (t = −3.1, p = 0.003, Fig. 5e and Table 1), suggesting a decrease over time in the CTIw at high elevations but not low elevations (Fig. 5e). When using the unweighted CTI (CTIuw), results were qualitatively the same for Forillon and Gatineau. At Mont-Mégantic, however, we found no effect of time and a clear and significant decrease in CTIuw with elevation for both the original and recent surveys (see Appendix S7).

**Figure 5.**
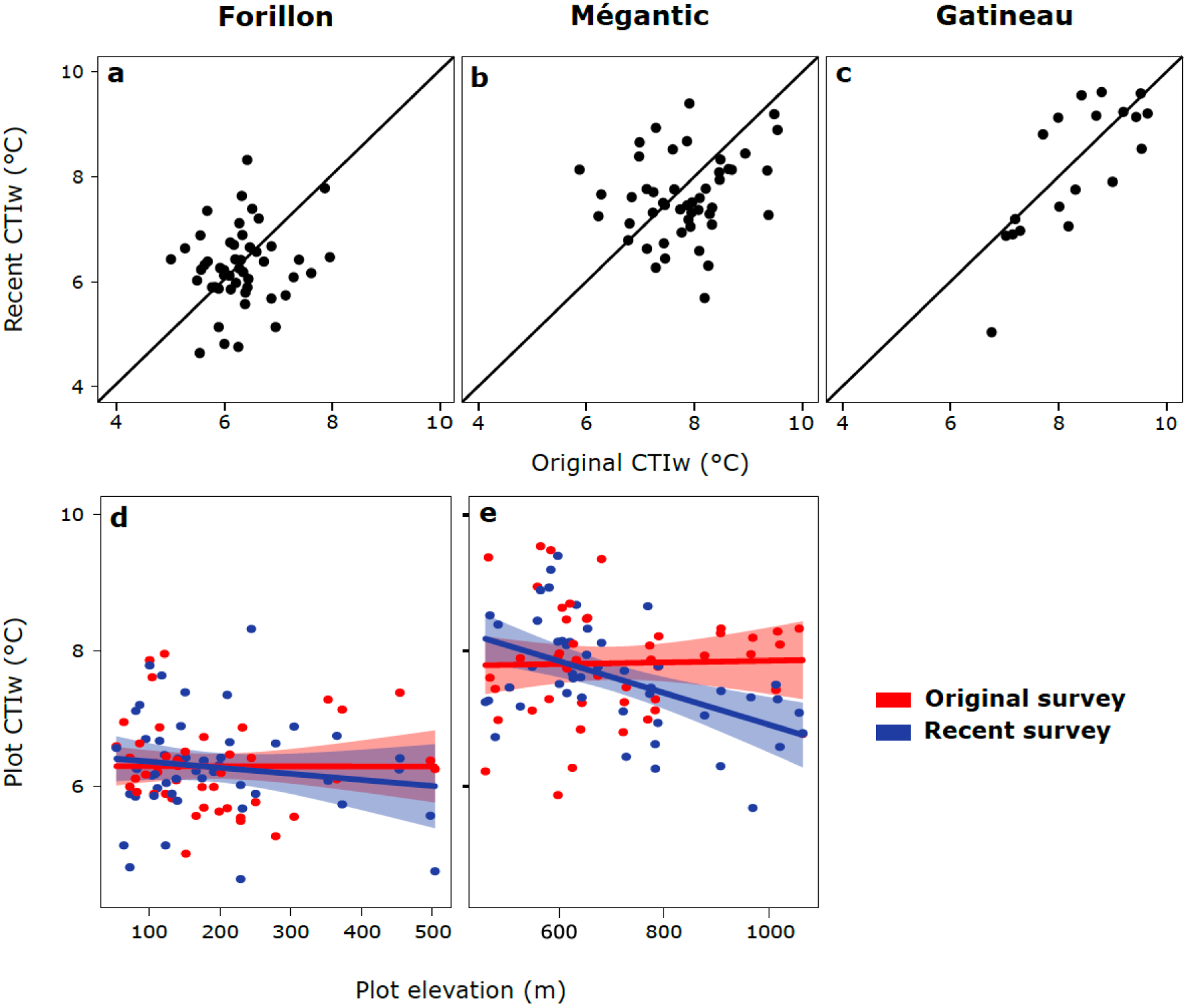
Community Temperature Indices (CTIw) during the two-time periods and across the elevational gradient. (a-c) Abundance-weighted indices (CTIw) at Forillon, Mont-Mégantic, and Gatineau, with the 1:1 line indicating no temporal change between two times. (d-e) Relationships between CTIw and elevation for each time period at Forillon and Mont-Mégantic. Red and blue illustrate original and recent surveys, respectively. Each point is a plot in all panels.

## Discussion

Many studies at single sites have revealed temporal changes in species distributions, community composition, or phenology that are consistent with predictions based on climate warming (Lenoir *et al.*, 2009; Bertrand *et al.*, 2011; Bernhardt-Römermann *et al.*, 2015; Sproull *et al.*, 2015; Ash *et al.*, 2017; Rogora *et al.*, 2018). However, with observational data (i.e., most long-term studies) it is always difficult to rule out alternative causes of temporal community change, such that comparative multi-site studies are needed to strengthen tests of the general hypothesis that biotic change over time has been influenced by climate warming (Verheyen *et al.*, 2017). In this study, we have taken advantage of a natural gradient in the degree of climate warming and of a protected area network in eastern Canada, combining three re-survey efforts totaling >100 plots to test whether greater warming has led to more marked changes in species distributions and community properties. Results were mostly consistent with our predictions, with the magnitude of biotic changes (i.e. elevational distributions, species richness, composition) most often increasing from Forillon Park in eastern Québec, where the warming trend has been relatively weak, to Mont-Mégantic where warming has been moderate, to Gatineau Park in western Québec where the warming trend has been the strongest. Results for community temperature indices were difficult to interpret, as discussed further below.

### Species’ elevational distributions

As predicted, species’ mean elevations shifted upward at Mont-Mégantic but not Forillon. There is no elevational gradient in Gatineau Park. On average, species at Mont-Mégantic moved toward higher elevations, as predicted if species are at least partially spatially tracking their temperature optima in response to warming (Kelly & Goulden, 2008; Savage & Vellend, 2015; Sproull *et al.*, 2015).

The rate of elevational shift for the understorey plants at Mont-Mégantic (~10 m.decade^−1^) is close to the global average of 11 m.decade^−1^ reported in the meta-analysis of Chen *et al.*, 2011), although individual studies have reported higher values (e.g., ~22 m.decade^−1^ in southern California; Kelly & Goulden, 2008) and lower values (e.g., no shift in elevation in Montana; Klasner & Fagre, 2002). However, direct comparison among studies in different regions is complicated by different degrees of warming over the relevant time frames in different places. Moreover, there has been relatively few studies in North-America, making our study not only a novel general contribution to global change biology, but also a valuable regional-scale contribution to our knowledge of changes in species distributions along elevation gradients in eastern North-America.

Although the gradients in Forillon and Mont-Mégantic cover similar elevational ranges (~500-600 m), the vegetation gradient is less pronounced in Forillon Park than at Mont-Mégantic. For instance, Forillon’s high elevation summits are not as predictably dominated by boreal forest as they are at Mont-Mégantic. This can be seen in the weaker relationships between plot richness and CTI with elevation at Forillon contrary to Mont-Mégantic (Figs. 3d-e, 5 d-e, Appendix S7). Despite these differences, the clear absence of any shift in elevational distributions in Forillon Park is consistent with the hypothesis that climate warming is the probable cause of elevational distribution shifts at Mont Megantic (and elsewhere).

### Species richness, composition, and heterogeneity

Since warm areas tend to have higher local plant diversity than cold areas, climate warming is predicted to increase local plant diversity in many regions (Vellend *et al.*, 2017). Consistent with our prediction, there was no significant temporal change in species richness over ~40 years at Forillon but significant increases were found at Mont-Mégantic and Gatineau. Some other studies in regions that have experienced warming have also found increases of local vascular plant diversity (Klanderud & Birks, 2003; Walther *et al.*, 2005; Stockli *et al.*, 2012; Steinbauer *et al.*, 2018), although temporal changes in species richness are highly variable (Verheyen *et al.*, 2012; Vellend *et al.*, 2013b).

We found significant temporal shifts in understorey community composition in all three parks, consistent with many studies in the literature showing species turnover through time (Magurran *et al.*, 2010; Dornelas *et al.*, 2014; Shi *et al.*, 2015). Comparisons among parks were consistent with our predictions, with the magnitude of community shifts (R^2^) following the gradient of warming: Forillon < Mont-Mégantic < Gatineau. However, we found no evidence of biotic homogenization, in contrast to many studies in literature (Jurasinski & Kreyling, 2007; Keith *et al.*, 2009; Zwiener *et al.*, 2018). In fact, our earlier study of Mont Mégantic reported significant biotic homogenization (Savage & Vellend 2015), and the difference with the present study appears to be largely due to differences in data processing and analysis. The raw community data were slightly different given our taxonomic standardization across surveys in different parks and a few differences in which woody plants were considered part of the understory vs. canopy (e.g., *Acer spicatum* was included in the understory in the current study but not the earlier one). More importantly,Savage & Vellend (2015) first used a fourth-root transformation of abundance data prior to calculating Bray-Curtis differences (a recommendation in the PRIMER software;Anderson *et al.*, 2008), whereas we saw no clear justification for this in the present study. Applying the same transformation to our data revealed significant biotic homogenization for Mont Mégantic, but not for the other two parks (results not shown). This is of negligible consequence for the present study, given that we did not have strong *a priori* predictions concerning beta diversity, although it is clear that the earlier result of biotic homogenization was not robust to alternative methods of analysis.

All observational studies involve uncertainty in making inferences about the cause of changes over space or time. Among potentially confounding factors that can underlie temporal community changes, succession is of potentially high importance. However, our study was designed specifically to minimize strong successional dynamics. We resurveyed plots originally surveyed in mature stands that have maintained closed canopies throughout the period of study. Importantly, we have no reason to suspect that forest dynamics (driven by factors other than climate) varies among our three parks in a way that aligns with the gradient of climate warming. As such, the best supported hypothesis for explaining the temporal changes we observed along the east-west gradient is that climate warming is a key driver.

Resurvey studies also raise questions about the comparability of surveys in different years and in different parks (Vellend et al. 2013a). In this study, in order to minimize differences between the six surveys, we paid close attention to taxonomic homogenization, and we consulted with botanists active in the 1970s (e.g., Colette Ansseau, a collaborator of M. Grandtner’s, and Z. Majcen) in order to reproduce the exact same field survey methods used in the original studies. One difference among parks we could not avoid was plot size, with smaller plots in Gatineau than in Forillon and Mégantic. It is predicted that in small communities, the importance of drift (stochastic changes in abundance) in driving community dynamics should be relatively high (Ricklefs & Lovette, 1999; Vellend, 2016). As such, all else being equal, one might have expected reduced detectability of deterministic community change over time in Gatineau, yet we found the opposite: a stronger temporal increase of α-diversity and a stronger directional shift in composition. Thus, if anything, we may have underestimated the difference between Gatineau and the other parks.

### Community temperature affinity (CTI)

The results for Community Temperature Indices (CTI) diverged most strongly from our predictions. Specifically, we failed to detect any temporal increase of CTI in Gatineau, and contrary to our prediction, we found a significant decrease of CTIw for high elevation plots at Mont-Mégantic (see Fig. 5e and Table 1). This result suggests a “cooling” in terms of community affinities to temperature at high elevation, which has actually been previously observed in the European Alps (Roth *et al.*, 2014). The fact that there was no such trend when using unweighted community temperature indices (CTIuw) indicates that changes in particular species’ abundances drove the result for CTIw.

In particular, two of the most abundant species experienced major temporal changes: (i) *Oxalis acetosella* L. (known also as *Oxalis montana* Raf.) decreased in average abundance and (ii) *Dryopteris carthusiana* (Vill.) H.P. Fuchs increased in abundance (see Appendix S8). *Oxalis acetosella* had a Species Temperature Index (STI) of 8.6 °C. This species was often found at unusually high abundance in the original surveys at Mont-Mégantic, especially at high elevation (>800 m). On average, *O. acetosella* contributed ~74% to CTIw values for high elevation plots in the original survey, while contributing only ~8% in the recent survey (see Appendix S8). Given abundance reductions at high elevation, the abundance-weighted elevation of this species declined more than any other, which represents an exception among the full set of species (*O. acetosella* is the right-most point in Fig. 2b), but which has a major effect on CTIw values. In contrast, *Dryopteris carthusiana* (STI = 7.6 °C) was not particularly abundant at high elevation in the original surveys but became very abundant in the recent surveys. The contribution of *D. carthusiana* to CTIw for plots at high elevation (>800 m) increased from 9.5% to 47%. At Mont-Mégantic, *O. acetosella* is more strongly associated with high elevation forests (i.e., colder sites) than is *D. carthusiana*, and so their changes in abundance are in one sense consistent with the hypothesis that warming is a major driver of vegetation change. But since the estimated STI (using independent data) was actually higher for *O. acetosella* than *D. carthusiana*, the changes in abundance caused a decline in high-elevation CTIw. In sum, the high sensitivity of CTI to the dynamics of individual species, combined with uncertainty in STI values (see also below), may reduce the degree to which CTI acts as an indicator of climate warming.

The calculation and interpretation of CTI has several limitations. First, Species Temperature Indices (STI) are calculated based on recorded species occurrences, but for many species we have limited knowledge of geographic distributions, especially in northern regions or at high elevation. Second, the assumption that median temperature represents a species’ optimum is unverified (Rodriguez-Sanchez *et al.*, 2012). As mentioned above, STI is greater (warmer) for *Oxalis* than for *Dryopteris* due to the more northern distribution of *Dryopteris.* However, in eastern North America *Oxalis* is known to be more abundant in coniferous forests at high elevation while *Dryopteris* is more widely distributed along elevation gradient. Thus, if we used data from occurrences along elevational gradients (i.e., at Mont-Mégantic), *Oxalis* would have a *lower* STI than *Dryopteris*. In other studies, CTI has been shown to increase as predicted by warming (Devictor *et al.*, 2008; Lindström *et al.*, 2012; Bowler *et al.*, 2015). In our study system, STIs and therefore CTIs come with considerable uncertainty.

In sum, we have provided empirical evidence of vegetation changes in eastern Canada that are largely consistent with the east-west gradient in warming. Explicit comparisons of community change among regions with variable climatic histories appears to be a powerful method for increasing the confidence with which biotic trends can be attributed to climate warming. Many unknowns remain, such as the functional attributes of “loser” and “winner” species, and the extent to which adaptive changes within species might also contribute to warming responses. Continuing to exploit historical data sources of all kinds can help advance global change science.

## Acknowledgements

We thank the field and lab assistants who contributed to this project: Diane Auberson-Lavoie, Melissa Paquette, and Sara Gaignard. This work was made possible thanks to the support of park managers, especially Camille-Antoine Ouimet (Mont-Mégantic) and Daniel Sigouin (Forillon). We also thank Guillaume Blanchet, Raphael Aussenac and Joanie Van De Walle for valuable input on various aspects of this project, in particular statistical analysis. Finally, thanks to Pauline Palmas and Arnault Lalanne for providing comments on an earlier version of the manuscript. Funding was provided by the Natural Sciences and Engineering Research Council, Canada.

## Appendices

- Appendix S1 - Climatic trends in three regions of Québec, Canada
- Appendix S2 - Taxonomic standardization between surveys
- Appendix S3 - Species Temperature Index (STI) database
- Appendix S4 - Mean abundance-weighted elevation and number of occurrences per species per survey in Forillon and Mont-Mégantic
- Appendix S5 - Species occurrences per survey (number of plots where species were recorded)
- Appendix S6 - Global ordination of community composition for all three parks in both time-periods
- Appendix S7 - Results for unweighted Community Temperature Indices (CTIuw)
- Appendix S8 - Individual species contributions to Community Temperature Indices (CTIw) for high elevation plots at Mont-Mégantic

Contribution
ABS and MV designed the study and wrote the manuscript collaboratively (with ABS as leader); ABS collected and analyzed the data; SV extracted climatic and species distribution data and provided input on the manuscript.

